# Central Glucagon-Like Peptide-1 Agonism Ameliorates Colonic Inflammation in a Murine Model of Inflammatory Bowel Disease

**DOI:** 10.64898/2026.10.22.761061

**Authors:** Biren Desai, Kendra L. Francis, Caeley L. Bryan, Jessie Connolly, Thanushree Karunagaran, Sara Mathan, Karina Sharma, Vincent Damian, Christina A. Watts, Pique P. Choi, Kelly Kadlec, Bao Anh Phan, Maria C. Pacheco, Kimberly M. Alonge, Gregory J. Morton, Michael W. Schwartz, Jarrad M. Scarlett

## Abstract

Glucagon-like peptide-1 receptor agonists (GLP-1RAs) have revolutionized the treatment of obesity and type 2 diabetes (T2D). These drugs also have anti-inflammatory effects that have recently been extended to the treatment of inflammatory bowel disease (IBD). Whether the mechanism underlying this benefit involves a direct action on the intestine or instead is mediated via GLP-1Rs in the brain, however, is unknown. Based on evidence implicating the brain as the target for the anti-obesity and anti-inflammatory effects of GLP-1RAs, we sought to determine if central GLP-1R signaling is sufficient to explain its intestinal anti-inflammatory effects. To this end, we compared the efficacy of peripheral vs central administration of the GLP-1RA, semaglutide (SEMA), in diet-induced obese (DIO) mice with dextran sodium sulfate (DSS)-induced colitis. We report that compared to vehicle, both colitis severity and histopathological injury following DSS administration are reduced by peripheral administration of SEMA. We also report that this benefit is largely recapitulated by intracerebroventricular administration of SEMA at a low dose that, when administered peripherally, is below that needed to affect food intake and body weight. We conclude that in a mouse model of colitis, activation of GLP-1Rs in the brain is sufficient to explain the intestinal anti-inflammatory effect conferred by GLP-1R agonism.

## INTRODUCTION

Inflammatory bowel disease (IBD), comprising ulcerative colitis (UC) and Crohn’s disease (CD), is a chronic inflammatory disorder of the gastrointestinal (GI) tract that affects approximately 7 million people worldwide [1]. Neither condition has an established medical cure, and both contribute substantially to disability, reduced quality of life, and health care expenditures [2–4]. IBD susceptibility is influenced by genetic predisposition, immune dysregulation, gut microbiome composition, and environmental exposures, including diet, and while biologic therapies and small-molecule inhibitors have expanded treatment options [5–7], many patients do not achieve sustained remission. Moreover, existing immunosuppressive therapies increase the risk of infection, malignancy, and end-organ toxicity [8–10]. These limitations highlight the need for therapeutic advances that attenuate intestinal inflammation without need for immunosuppression.

Obesity and type 2 diabetes (T2D) are increasingly prevalent comorbidities in patients with IBD, a trend of particular concern given the rising incidence of these metabolic disorders among children and adolescents [11, 12]. Both conditions are also associated with increased rates of IBD-related hospitalization, disease flares, and need for surgery [8, 13]. Incretin-based therapies, including glucagon-like peptide-1 receptor agonists (GLP-1RAs), represent major advances in obesity and T2D treatment [9], and these drugs have also been linked to improved IBD outcomes [10, 14, 15]. Specifically, GLP-1RA treatment associates with higher rates of IBD remission [16] and reduced rates of IBD-related hospitalization, surgery, and corticosteroid use in this patient population [17]. What remains unclear is how these salutary effects are mediated.

In addition to reducing body weight and improving glycemic control [18], GLP1-RAs are also reported to have anti-inflammatory effects independent of weight loss [19]. In the gastrointestinal tract, peripheral administration of GLP-1RAs suppresses inflammatory signaling and cytokine production [20], modulates mucosal T-cell responses [21], enhances intestinal barrier integrity and mucus production [20, 22] and alters host-microbiome interactions [21]. While a peripheral mechanism of action is often invoked to explain these effects, GLP-1R activation in the brain is implicated in the anti-inflammatory actions of these drugs. For example, central administration of GLP-1RAs can attenuate both lipopolysaccharide-induced inflammation and sepsis-associated pathology in a cecal slurry model of polymicrobial sepsis in mice [19, 23]. In the setting of IBD, therefore, a central mechanism could potentially account for the benefit conferred by GLP-1RAs.

In the current work, we sought to determine if central GLP-1R activation is sufficient to attenuate DSS-induced colitis in mice with diet-induced obesity (DIO). We first demonstrated that in this model, intestinal inflammation and colitis severity are significantly reduced by chronic peripheral SEMA infusion. We then showed that intestinal inflammation and disease severity are similarly attenuated when SEMA is administered by intracerebroventricular (icv) infusion. Importantly, this effect occurs at a dose 10-fold below that needed for benefit when SEMA is administered subcutaneously (sc) – a dose that when administered systemically, has no detectable effect on food intake or body weight. Collectively, these findings demonstrate that central SEMA administration is sufficient to attenuate experimental colitis in DIO mice and support a potential role for CNS GLP-1R signaling in the intestinal anti-inflammatory effects of GLP-1R agonism.

## RESULTS

### Peripheral SEMA administration attenuates IBD clinical disease activity in chow-fed mice with DSS-induced colitis

To establish proof of principle for our drug-delivery approach, we examined whether continuous peripheral infusion of SEMA via osmotic minipump could recapitulate the attenuation of DSS-induced colitis previously reported with daily subcutaneous (sc) injections in chow-fed mice (Figures S1A-B) [20, 24]. Chow-fed, male mice underwent implantation of a subcutaneous osmotic minipump containing SEMA (40μg/kg/day) or vehicle control (VEH) (Figure S2A). As expected, SEMA reduced body weight over the initial 15-day treatment period (mean weight loss –5.48% ± 1.28%, p=<0.001 vs. VEH) (Figure S2B). After body weight stabilized, mice received 2% DSS or untreated drinking water for 10 days while SEMA or VEH infusion continued, generating four groups: VEH/VEH, SEMA/VEH, VEH/DSS, and SEMA/DSS (Figure S2A). VEH-treated mice were pair-watered to their SEMA-treated counterparts to ensure matched DSS intake across groups [25]. Since weight loss is a known effect of SEMA, this variable was assessed separately from clinical IBD disease activity index (DAI) scoring, and weight loss during DSS exposure tended to be lower in SEMA/DSS-treated mice than in controls (VEH/DSS-treated mice) (–11.34% vs. –14.72%, p=0.087, Figure S2C).

Clinical disease activity was assessed using a modified DAI comprising stool consistency and rectal bleeding scores, each ranging from 0 to 4, for a total score ranging from 0–8. As expected, DAI scores were elevated in VEH/DSS mice compared to control groups that were not exposed to DSS. These clinical abnormalities were significantly attenuated in SEMA/DSS-treated mice beginning on Day 4 and continuing through sacrifice on Day 10 (Figure S2D). At study conclusion, stool consistency and rectal bleeding scores were significantly improved in SEMA/DSS-treated mice compared to VEH/DSS-treated mice (Figure S2E, F). Consistent with the improvement in clinical disease activity, SEMA treatment also attenuated the DSS-associated increase in spleen weight, a marker of systemic inflammation [26] (Figure S3A). Colon length, which decreases with increasing inflammation and colitis severity [27], was significantly reduced in VEH/DSS-treated mice relative to both SEMA/DSS– and VEH/VEH-treated mice (Figure S3B). Together, these findings demonstrate that in chow-fed mice with DSS-induced colitis, clinical disease activity, splenic enlargement, and colon shortening were each attenuated by continuous sc SEMA infusion.

### Peripheral SEMA administration attenuates IBD clinical disease activity in DIO mice with DSS-induced colitis

Since DIO both exacerbates DSS-induced colitis severity and associates with increased responsiveness to GLP-1RA drugs, we next examined the effect of continuous sc SEMA infusion on DSS-induced colitis in mice with DIO. Mice were maintained on a 60% high-fat diet (HFD) for 6 weeks before treatment and remained on this diet throughout the study. SEMA (40 μg/kg/day) or VEH was administered through subcutaneously implanted osmotic minipumps using the protocol described above (Figure 1A). As expected, SEMA induced a rapid, robust and sustained reduction of body weight of approximately 18% over the first 12 days, after which body weight stabilized (Figure 1B). Before DSS exposure, mean body weight was 41.56 ± 1.27 g in VEH-treated mice and 31.93 ± 0.95 g in SEMA-treated mice (*P* < 0.0001; Figure 1B). After body weight stabilization, mice received 2% DSS or untreated drinking water for 10 days while sc SEMA or VEH infusion continued (Figure 1A). As in the chow-fed cohort, VEH/DSS mice were pair-watered to SEMA/DSS mice to control for potential differences in DSS exposure (Figure S4A) [25]. Body weight change during DSS exposure was analyzed separately from the DAI score and did not differ between VEH/DSS and SEMA/DSS mice (VEH/DSS, –7.37% ± 1.10% vs. SEMA/DSS, – 7.92% ± 1.25%; *P* = 0.78; Figure 1C).

**Figure 1.**
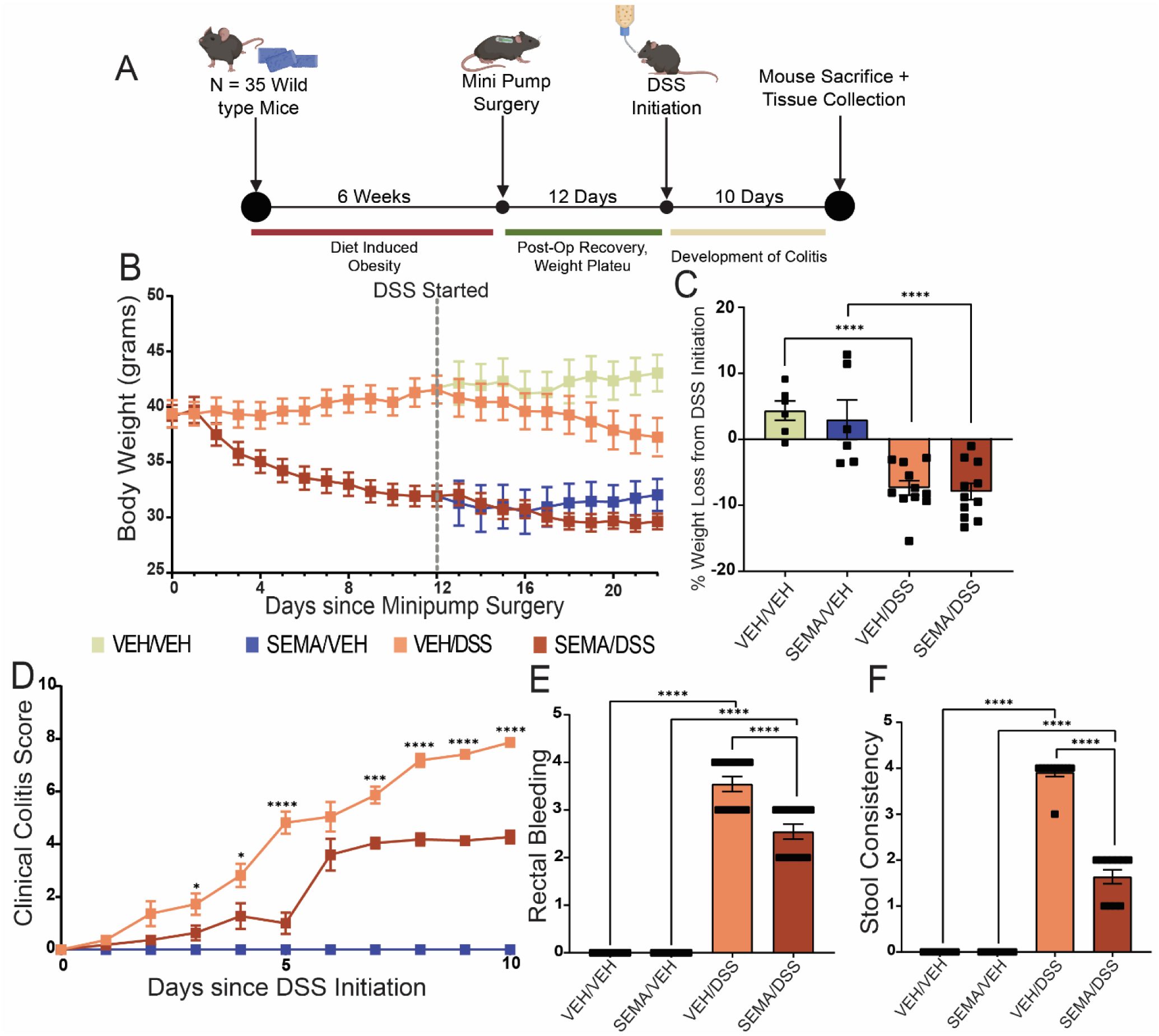
Peripheral SEMA improves clinical outcomes in DSS-treated DIO mice. (**A**) Experimental timeline, (**B**) body weight, (**C**) total percent weight loss during DSS-treatment, (**D**) clinical colitis scores, (**E**) rectal bleeding, and (**F**) stool consistency in DIO mice that received continuous subcutaneous infusion of either SEMA (40 μg/kg/d) or its VEH and subsequently received either 2% DSS or VEH in the drinking water for the final 10 days. Water intake was paired between groups to ensure matched DSS exposure. *n* = 6-12 per group. Mean ± SEM. Non-parametric t-test comparing VEH/DSS and SEMA/DSS in (C). 2-way ANOVA followed by Fisher’s post hoc tests for (D-F). SEMA: Semaglutide; VEH: Vehicle; DSS: Dextran Sodium Sulfate; DIO: Diet induced obesity. ****P< 0.0001, ***P<

In DIO mice, DSS exposure increased DAI scores relative to controls not exposed to DSS. Beginning on Day 3, SEMA/DSS mice exhibited lower DAI scores than VEH/DSS mice, and this difference persisted throughout the study (Figure 1D). At study completion, mean DAI scores were reduced by 46% in mice receiving SEMA (7.86 ± 0.07 in VEH/DSS mice and 4.27 ± 0.25 in SEMA/DSS mice; *P* < 0.0001) (Figure 1D), reflected by marked reductions of both rectal bleeding and stool consistency scores (Figure 1E, F). Thus, continuous peripheral SEMA infusion attenuates clinical disease activity in both DIO and lean, chow-fed mice with DSS-induced colitis.

### Central SEMA administration attenuates IBD clinical disease activity in DIO mice with DSS-induced colitis

To determine whether central SEMA administration can reproduce the protective effects observed with peripheral treatment, DIO mice (maintained on an HFD for 8 wk) were implanted with osmotic minipumps for continuous intracerebroventricular (icv) infusion of either SEMA (4 μg/kg/day) or VEH (Figure 2A). The dose selected for central administration was 10-fold lower than the dose for subcutaneous infusion, and below that needed to reduce food intake or body weight when administered peripherally (Figure S5A, B). Before DSS exposure, icv SEMA reduced body weight by approximately 12%, with body weight plateauing by Day 12 (mean BW pre-DSS: icv Veh 34.34 ± 0.69g vs. icv SEMA 30.67 ± 0.85g; *P* = <0.01) (Figure 2B). Mice subsequently received 2% DSS or untreated drinking water for 10 days while icv infusion continued (Figure 2A). VEH-treated groups were pair-watered to their corresponding SEMA-treated groups to control for potential differences in water or DSS intake (Figure S4B) [25]. Body weight loss during DSS exposure was not included in DAI score but did not differ significantly between groups (icv VEH/DSS, –3.38% ± 0.86% vs. icv SEMA/DSS, –6.02% ± 1.87%; P = 0.2384; Figure 2C).

**Figure 2.**
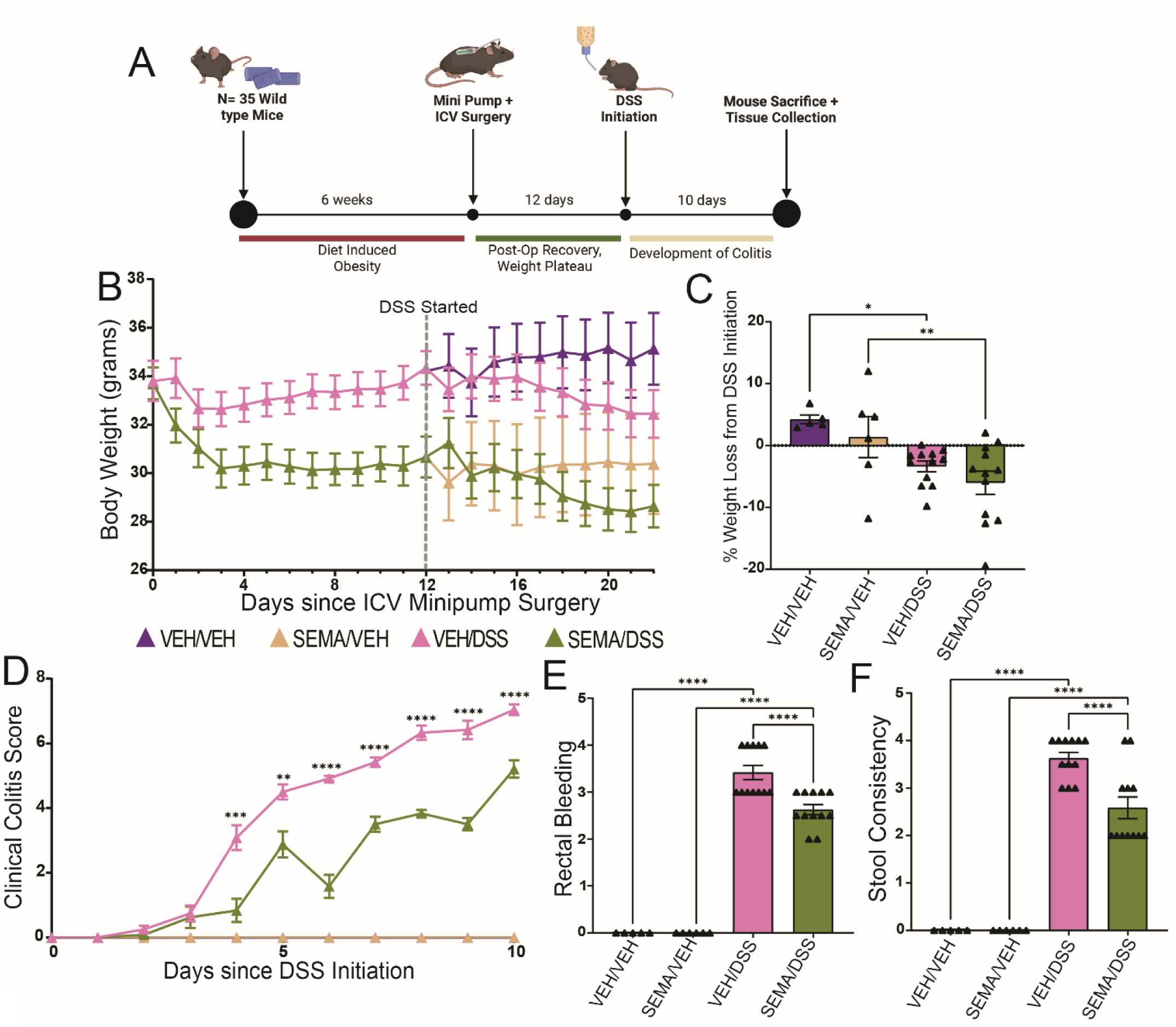
Central SEMA administration improves clinical outcomes in DSS-exposed DIO mice. (**A**) Experimental timeline, (**B**) body weight, (**C**) total percent weight loss during DSS, (**D**) clinical colitis scores, (**E**) rectal bleeding score, and (**F**) stool consistency, in DIO mice that received continuous icv infusion of either SEMA (4 μg/kg/d) or its VEH and subsequently received either 2% DSS or VEH in the drinking water for that last 10 days. *n* = 6-12 per group. Mean ± SEM. Non-parametric t-test comparing VEH/DSS and SEMA/DSS in (C). 2-way ANOVA followed by Fisher’s post hoc tests for (D-G). SEMA: Semaglutide; VEH: Vehicle; DSS: Dextran Sodium Sulfate; DIO: Diet induced obesity; ICV: Intracerebroventricular. ****P< 0.0001, ***P< 0.001, **P< 0.01, *P< 0.05

As was observed during sc SEMA administration, the effect of DSS exposure to increase DAI scores (relative to controls not exposed to DSS) was blunted by icv SEMA treatment (Figure 2D). This effect became detectable beginning on Day 4 and persisted throughout the study, with DAI scores being 26% lower than icv VEH/DSS mice (icv VEH/DSS, 7.04 ± 0.17 vs. icv SEMA/DSS, 5.21 ± 0.27; P < 0.0001; Figure 2D). This reduction was accompanied by lower rectal bleeding and stool consistency scores (Figure 2E, F), demonstrating that continuous central SEMA infusion is sufficient to mimic the effect of systemic SEMA to attenuate clinical disease activity in DIO mice with DSS-induced colitis.

### Peripheral and central SEMA protect against DSS-associated histopathologic damage in DIO mice

We next assessed gross pathological and histopathological outcomes following peripheral and central SEMA administration in DIO mice with DSS-induced colitis. In the peripheral treatment cohort, spleen weight was greater in VEH/DSS mice than in VEH/VEH controls, consistent with increased systemic inflammation, whereas spleen weights in SEMA/DSS mice were comparable to those in VEH/VEH controls (Figure 3A). Peripheral SEMA also attenuated DSS-induced colon shortening: colons were significantly longer in SEMA/DSS mice than in VEH/DSS mice (8.19 ± 0.15 vs. 6.85 ± 0.19 cm; P < 0.0001; Figure 3B). Unexpectedly, SEMA also increased colon length in mice not exposed to DSS (SEMA/VEH, 9.18 ± 0.35 cm vs. VEH/VEH, 7.88 ± 0.12 cm; P = 0.0007; Figure 3B), indicating that this effect was not restricted to DSS-treated mice. Histopathological analysis of hematoxylin and eosin–stained colon sections revealed higher histological damage scores in VEH/DSS mice than in VEH/VEH controls (Figure 3C, D), and this effect was also blunted by continuous sc SEMA administration (SEMA/DSS, 7.17 ± 0.30 vs. VEH/DSS, 9.09 ± 0.53; P = 0.0004; Figure 3C, D).

**Figure 3.**
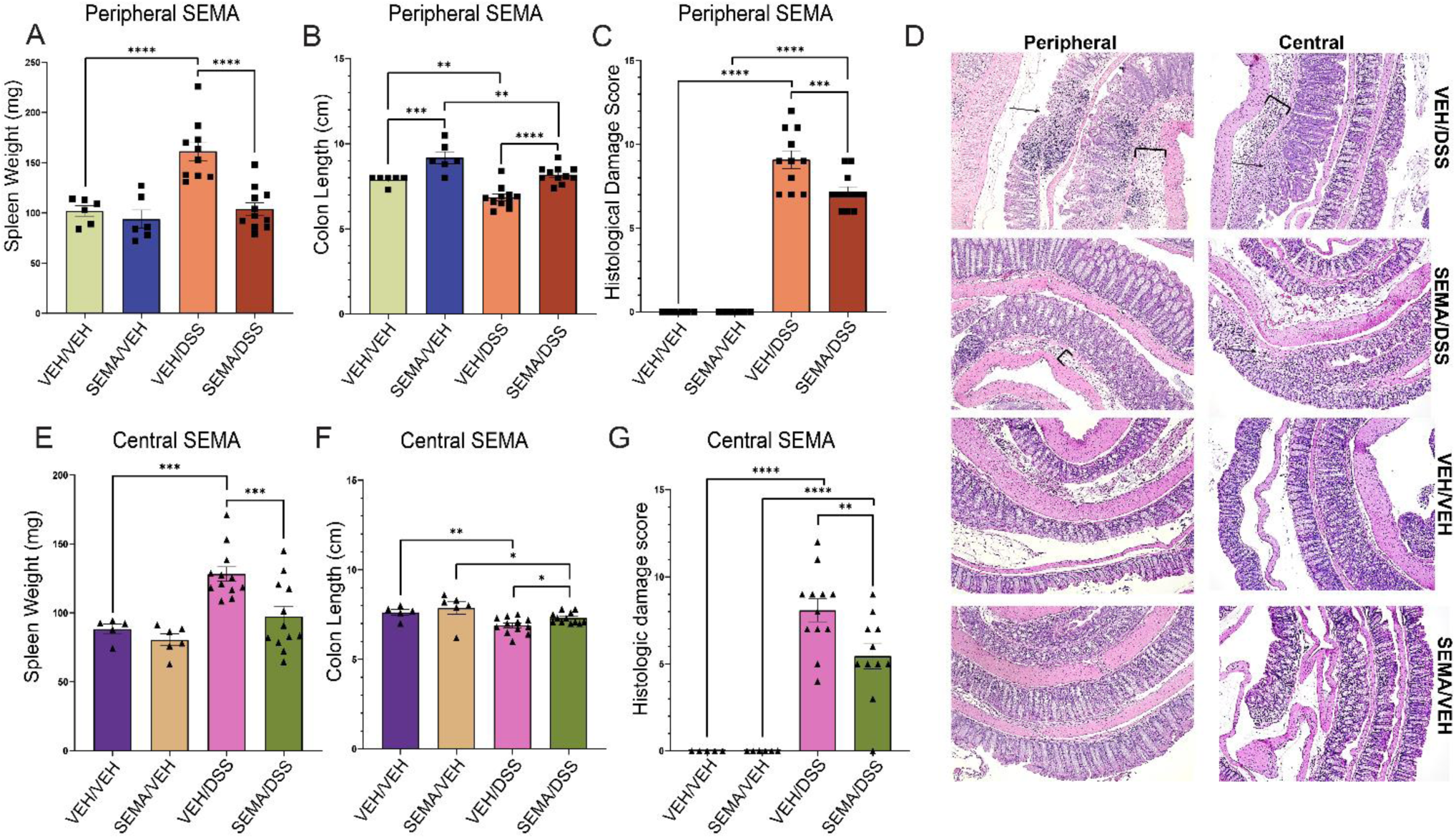
Peripheral and central SEMA administration each improve both pathologic and histopathologic outcomes in DSS-treated DIO. (A, E) Spleen weights, (B, F) colon length, (C, G) histopathologic damage scores and (D) representative H&E images in DIO mice exposed to either 2 % DSS or VEH in the drinking water and that received either peripheral (40 μg/kg/d) or central (4 μg/kg/d) administration of SEMA, respectively, relative to VEH-control. Inflammatory cell infiltrate shown in (**D)** causing *1)* expansion of the submucosal layer shown by brackets and *2)* epithelial damage disrupting normal crypt architecture shown by arrows. *n* = 11 – 12 for VEH/DSS and Sema/DSS groups, *n* = 5 – 6 for SEMA/VEH and VEH/VEH groups. 2-way ANOVA followed by Fisher’s post hoc tests. SEMA: Semaglutide; VEH: Vehicle; DSS: Dextran Sodium Sulfate; DIO: Diet induced obesity; H&E: Hematoxylin and Eosin. ****P< 0.0001, ***P< 0.001, **P< 0.01, *P< 0.05.

Following icv administration, SEMA similarly attenuated the DSS-associated increase in spleen weight (Figure 3E), and colon length was slightly but significantly greater in icv SEMA/DSS mice than in icv VEH/DSS mice (7.31 ± 0.09 vs. 6.89 ± 0.13 cm; P < 0.0001; Figure 3F). Increased histological damage scores in icv VEH/DSS mice (relative to icv VEH/VEH controls) were also reduced by central SEMA treatment (icv SEMA/DSS, 5.46 ± 0.74 vs. icv VEH/DSS, 8.08 ± 0.66; P = 0.0034; Figure 3D, G). Unlike systemic SEMA, however, central SEMA administration did not increase colon length in mice that did not receive DSS. Together, these findings demonstrate that whether administered peripherally or centrally at a much lower dose, SEMA attenuates clinical and histochemical manifestations of DSS-induced colitis with similar efficacy.

### Peripheral and central SEMA differentially modulate intestinal junction proteins in DIO animals

In the colon, claudin-1 and E-cadherin are components of epithelial tight and adherens junctions, respectively. The reduced abundance of each of these proteins in mice with DSS-induced colitis and other intestinal inflammatory diseases is suggestive of impaired intestinal barrier integrity [28–31]. To assess the effect of SEMA on these biomarkers, colonic sections were immunostained for claudin-1 and E-cadherin, and mean fluorescence intensity was quantified. In the peripheral treatment cohort, DSS reduced staining for both proteins in VEH-treated mice relative to VEH/VEH controls, as expected, and in mice not exposed to DSS, sc SEMA infusion had no significant effect on either biomarker (Figure 4A-C). In DSS-treated mice, however, peripheral SEMA increased staining for both claudin-1 (SEMA/DSS, 67.11% ± 2.30% vs. VEH/DSS, 54.72% ± 2.22%; P = 0.002) and E-cadherin (SEMA/DSS, 62.04% ± 3.27% vs. VEH/DSS, 37.71% ± 3.99%; P < 0.0001) relative to sc vehicle infusion. Indeed, systemic SEMA administration restored staining for both proteins to values comparable to VEH/VEH controls (Figure 4A–C).

**Figure 4:**
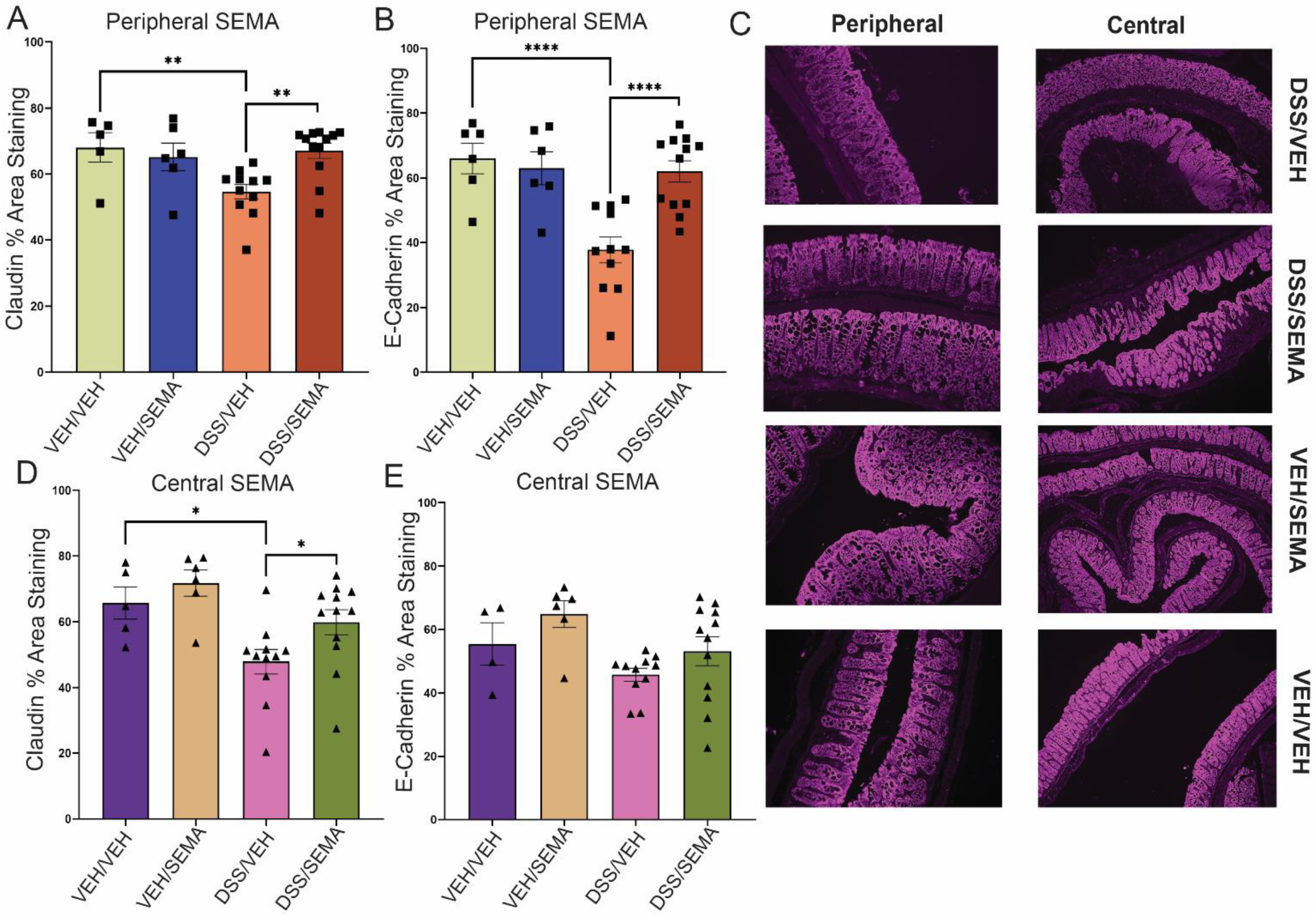
Effect of peripheral and central SEMA administration on the colonic mucosal barrier in DSS-treated DIO mice. (A, D) Claudin, (B, E) E-Cadherin percent area staining and (**C**) representative images of Claudin in DIO mice exposed to either 2 % DSS or VEH in the drinking water and that received either peripheral (40 μg/kg/d) or central (4 μg/kg/d) administration of SEMA, respectively, relative to VEH-control. Increased fluorescence is seen in the DSS/SEMA group compared to the DSS/VEH group. *n =* 6 – 12/group. 2-way ANOVA with post-hoc Fisher’s testing for (A-B, D-E). DSS: Dextran Sodium Sulfate; VEH: Vehicle; IHC: Immunohistochemistry. ****P< 0.0001, ***P< 0.001, **P< 0.01, *P< 0.05.

In the icv treatment cohorts, claudin-1 staining was reduced in icv VEH/DSS mice relative to icv VEH/VEH controls, as expected, and icv SEMA increased claudin-1 staining in DSS-treated mice (icv SEMA/DSS, 59.86% ± 3.80% vs. icv VEH/DSS, 47.89% ± 3.74%; P = 0.025; Figure 4C, D). In contrast, E-cadherin staining in icv SEMA/DSS mice did not differ significantly from that in either icv VEH/DSS mice or icv VEH/VEH controls (Figure 4E). Thus, peripheral SEMA attenuates DSS-associated reductions in both claudin-1 and E-cadherin staining, whereas central SEMA attenuates the decrease of claudin-1 staining without a statistically detectable effects on E-cadherin content.

### Peripheral and central SEMA decreased DSS-associated inflammatory components of the extracellular matrix

The intestinal extracellular matrix (ECM) represents another axis governing the interplay between luminal contents, the immune system, and barrier function. Based on previously reported pro-inflammatory changes in chondroitin sulfate glycosaminoglycan (CS-GAG) composition during colitis (CS-GAGs) [32–34] including an increase in the abundance of the di-sulfated proinflammatory CS isomer, CS-E (4S6S sulfation pattern) [35], we quantified the relative abundance of colonic CS-GAG disaccharide species by enzymatic digestion followed by liquid chromatography–tandem mass spectrometry (Figure 5A; Supplementary Table 1). As expected, intestinal CS-E content was increased in DSS-treated mice, and this effect was significantly reduced by peripheral SEMA administration, while having no effect in mice not exposed to DSS (Figure 5B, C; Supplementary Table 1). Central SEMA administration similarly reduced the relative abundance of CS-E in DSS-treated mice compared with icv VEH/DSS controls (Figure 5D, E; Supplementary Table 1). Further, CS-E relative abundance was positively correlated with histological inflammation scores in both the peripheral (P = 0.004; R² = 0.33) and central treatment cohorts (P = 0.015; R² = 0.24). Whether given peripherally or centrally, therefore, SEMA attenuates the effect of DSS to increase the relative abundance of the colonic proinflammatory CS-E isomer, and the magnitude of this effect is proportional to the reduction of histological inflammation. These results identify a potential association between ECM remodeling and the protective effects of peripheral and central GLP-1R signaling.

**Figure 5:**
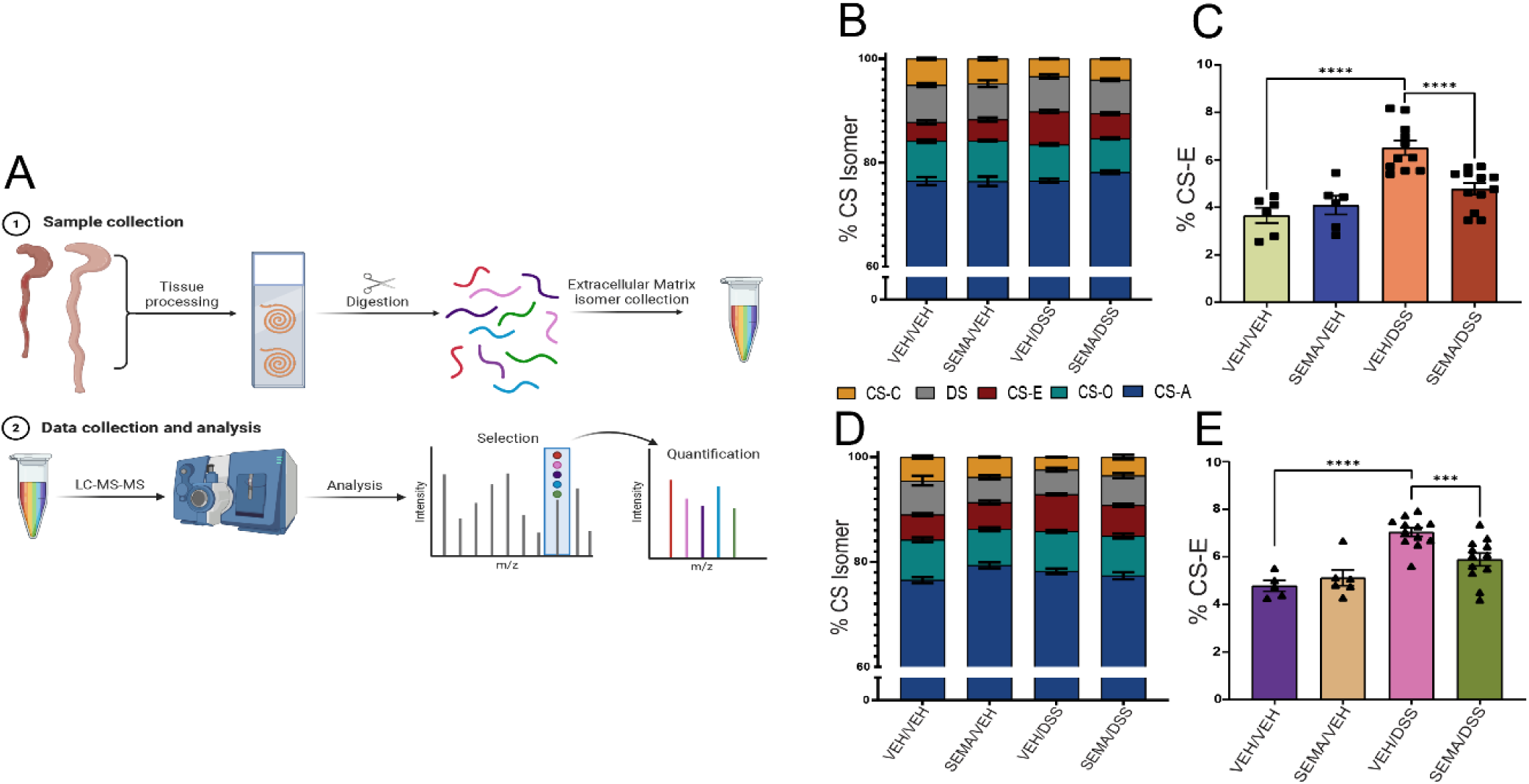
Peripheral and central administration of SEMA decreases the abundance of the pro-inflammatory colonic CS-E chondroitin sulfate in DSS –treated DIO mice. (A) Tissue processing steps for LC-MS/MS. Colonic (B, D) %CS isomer breakdown and (C, E) % CS-E analyzed using LC-MS/MS in DIO mice exposed to either 2 % DSS or VEH in the drinking water and that received either peripheral (40 μg/kg/d) or central (4 μg/kg/d) administration of SEMA, respectively, relative to VEH-control. *n =* 6 – 12/group. 2-way ANOVA with post-hoc Fisher’s testing for (C, E). DSS: Dextran Sodium Sulfate; VEH: Vehicle; LC-MS/MS: liquid chromatography-tandem mass spectrometry; CS: Chondroitin sulfate. ****P< 0.0001, ***P< 0.001, **P< 0.01, *P< 0.05.

## DISCUSSION

GLP-1RAs are now established treatments for obesity and T2D, and emerging clinical evidence suggests that therapeutic benefit extends to patients with IBD [10, 14, 16, 17]. Beyond their metabolic effects, GLP-1RAs suppress systemic and tissue inflammation, partly through CNS GLP-1R signaling pathways [23], but whether the brain contributes to the salutary effects of these drugs in IBD is unknown. Here, we show that continuous peripheral SEMA administration reduces clinical disease activity and colonic injury in DIO mice with DSS-induced colitis, consistent with previous reports [20, 24], and that these intestinal anti-inflammatory effects are largely replicated by central administration of SEMA at a much lower dose. These findings indicate that central GLP-1R signaling reduces intestinal inflammation in a mouse model of colitis and implicates the brain as a target for the therapeutic benefit conferred by incretin therapy for patients with IBD.

The beneficial effects of peripheral SEMA administration extended to virtually all measured endpoints of DSS-induced colitis. These include splenic enlargement and colon shortening, histopathological evidence of intestinal injury, relative abundance of epithelial junctional proteins, and pro-inflammatory alteration of intestinal ECM components. A key finding is that these effects were largely recapitulated following intracerebroventricular (icv) SEMA infusion at a 10-fold lower dose – a dose that lacks metabolic efficacy when administered peripherally. While these findings therefore suggest that the protective effect of SEMA involves a central mechanism, our findings do not address whether SEMA also has direct anti-inflammatory effects on the intestine. Similarly, our studies do not address whether neuronal GLP1R signaling is *required* for the effect of SEMA to reduce intestinal inflammation, a question that we are currently pursuing.

The observed intestinal anti-inflammatory effects of central SEMA are consistent with previous evidence implicating CNS GLP-1Rs in the suppression of systemic inflammation [23]. Our findings extend this framework to experimental colitis and raise the possibility that brain-gut pathways modify the intestinal response to injury, although the relevant GLP-1R–expressing neuronal populations and downstream pathways remain to be identified. Studies using neuronal or tissue-specific GLP-1R activation and/or deletion will help to identify the neurons involved and clarify whether peripheral and central actions of SEMA converge on common intestinal targets.

An important consideration is the potential contribution of changes in body weight to the observed effects of SEMA. Both peripheral and central administration of SEMA resulted in significant weight loss in DIO mice prior to DSS initiation. To limit the contribution of ongoing SEMA-induced weight loss during colitis induction, DSS treatment was initiated only after body weight had stabilized at a plateau value, thus excluding the possibility that improvements in disease activity were the result of ongoing weight loss. Nevertheless, our findings do not establish weight-independent protection, as SEMA-treated mice entered the DSS treatment period at a lower body weight. Pair-feeding studies and comparisons with weight-matched controls would help clarify the contribution of preceding changes in food intake and body weight.

An interesting and unexpected finding is that whereas DSS-associated colon shortening was attenuated by SEMA regardless of the route of administration, peripheral (but not central) SEMA increased colon length in DIO mice not exposed to DSS. However, peripheral SEMA had no effect on colon length in chow-fed mice not exposed to DSS, implicating that this result is dependent on dietary composition. Previous studies show that a HFD can cause colon shortening independent of colitis [36].Our findings suggest that SEMA may attenuate HFD-induced colon shortening through intestinotrophic effects that are independent of central mechanisms.

The intestinal barrier is an obvious potential target for the therapeutic effect of GLP-1RA activation. The integrity of the epithelial barrier is critical for maintaining intestinal homeostasis, and disruption of tight junctions and the protective mucin layer is implicated in the pathogenesis and progression of intestinal inflammation [37, 38]. In our study, peripheral SEMA administration increased the epithelial tight junction protein abundance in DSS-treated mice relative to VEH/DSS-treated controls. Central SEMA administration produced a more modest response, significantly increasing Claudin abundance, with a nonsignificant trend toward increased E-cadherin staining (p=0.1538). These findings suggest that both peripheral and central pathways may contribute to SEMA-associated changes in the intestinal barrier.

The intestinal ECM is another important component of the intestinal barrier [39, 40]. We recently reported that the relative abundance of CS-E, a pro-inflammatory ECM component, is elevated in the colon of humans with IBD [35]. Consistent with this observation, we report that CS-E isomer abundance is increased in DSS-treated mice, and it is interesting to note that this effect was significantly blunted by SEMA, whether given peripherally or centrally. This finding extends the anti-inflammatory effect of GLP-1R activation to the intestinal ECM, a novel finding. Also novel is the ability of central GLP-1RA administration to alter intestinal ECM structures. To our knowledge, this is the first evidence suggestive of a brain-gut signaling pathway that regulates the intestinal ECM microenvironment. Whether this effect reflects direct neural regulation of intestinal ECM remodeling or occurs secondarily to other metabolic, epithelial, vascular, or immune changes remains to be determined.

This study has several limitations. Although central SEMA administration attenuated experimental colitis, receptor-specific loss-of-function studies are needed to establish the requirement for CNS GLP-1R signaling and identify the relevant neurocircuitry. The contribution made by preceding weight loss also remains unresolved, and changes in junctional protein staining do not directly demonstrate improved barrier function. Finally, because SEMA treatment was initiated before DSS exposure, our data assesses protection against the development of colitis rather than treatment of established disease. The rapid and severe phenotype induced by DSS further limits the window for evaluating interventions after disease onset. Studies initiating SEMA after the emergence of clinical symptoms, including in models of spontaneous but gradually progressive colitis such as occurs in *IL-10*–deficient mice, have greater potential to address therapeutic efficacy once intestinal inflammation is established.

In conclusion, peripheral SEMA administration attenuates clinical disease activity, colonic injury, and pathological changes in epithelial junctional proteins and ECM composition in DIO mice with DSS-induced colitis. Central administration of SEMA at a substantially lower dose reproduces most these benefits, indicating that neuronal GLP1R signaling is sufficient to explain many of the intestinal actions of GLP-1R agonism. These findings provide a rationale for future studies defining central and peripheral mechanisms underlying SEMA-mediated protection and identifying future therapeutic targets for patients with IBD.

## METHODS

### Animal model and experiments

Eight-week-old male C57BL/6J wild type mice purchased from Jackson Laboratory (Stock no. 000664, Bar Harbor, ME) were housed at the University of Washington South Lake Union Vivarium facility under specific pathogen-free conditions in a temperature-controlled room with a 14:10 light/dark cycle. Mice were provided with ad libitum access to water and fed either standard laboratory chow (5053; 4.5% kcal fat, LabDiet, MO) or high fat (HFD) rodent diet (60% kcal from fat, D12492i, Research Diets, NJ), unless otherwise stated. All procedures were performed in accordance with NIH guidelines for the care and use of animals and were approved by the Institutional Animal Care and Use Committee at the University of Washington (Seattle, Washington).

### Subcutaneous Minipump Surgery

Adult male mice maintained on either chow or HFD were assigned to weight-matched groups and implanted subcutaneously with an osmotic minipump (ALZET Model 1004; DURECT Corporation, CA) containing either vehicle (Tris buffered saline with tween; ThermoFischer Scientific, MA) or the GLP-1RA, SEMA (Novo Nordisk, Copenhagen, Denmark) at a dose of 40μg/kg/day as previously described [41]. Before implantation, minipumps were incubated in sterile saline at 37°C for 24 hours to normalize pump flow rate according to manufacturer’s instructions. Animals received a peri-operative dose of buprenorphine hydrochloride (Reckitt Benckiser Pharmaceuticals, VA) for analgesia and were monitored daily for body weight and wound integrity throughout the study.

### Intracerebroventricular (icv) Cannula and Minipump Surgery

To enable continuous delivery of vehicle or SEMA directly into the brain, six-week HFD-fed mice underwent lateral ventricular cannulation (Alzet Model 0008851 Brain Infusion Kit 3, DURECT Corporation, CA) at stereotaxic coordinates 0.7 mm posterior to bregma; 1.3 mm lateral, and 2.0 mm below the skull surface [42]. During the same surgical session, animals were implanted sc with an osmotic minipump (ALZET Model 1004; DURECT Corporation, CA) that was connected to the lateral ventricle to enable icv infusion of either vehicle or SEMA (4 μg/kg/day) as previously described [43–45]. Mice received a peri-operative dose of buprenorphine hydrochloride (Reckitt Benckiser Pharmaceuticals, VA) for analgesia and post-operative recovery and were monitored daily throughout the study.

### Induction of Chemical Colitis

Approximately fourteen days after initiation of continuous sc or icv infusion of vehicle or SEMA (the precise timing was based on weight stabilization), mice received either 2% dextran sodium sulfate (DSS) (36-50 kDa; MP Biomedical, CA, United States) dissolved in autoclaved drinking water or autoclaved drinking water alone (VEH) for 10 days. During DSS treatment, mice were monitored daily for body weight and water consumption. To control for potential effects of SEMA to reduce water intake, VEH/DSS-treated mice were paired-watered to match the water (and DSS) consumption of SEMA/DSS-treated mice [46, 47]. Disease activity index (DAI) scores were calculated daily based on the sum of degree of rectal bleeding (score of 0-4) and stool consistency (score of 0-4) [27]. Body weight was analyzed separately from DAI scoring given the confounding effect of SEMA to cause weight loss prior to DSS administration [48].

### Tissue Collection

At study completion, on Day 10 following initiation of either DSS or VEH drinking water, mice were euthanized and spleen and colon collected. Spleen weight was recorded and normalized to body weight. The colon was excised and flushed with 0.1M phosphate-buffered saline (PBS) to remove intraluminal contents. The colon was measured from the beginning of the cecum to the end of the rectum. Two small sections of distal colon were cut and either flash frozen at –80°C or placed in RNAlater solution (Thermo Fisher Scientific, AM7030 Waltham, MA). The remaining colon was fixed in 10% zinc-buffered formalin (ZBF) for histological processing. Sections of spleen were stored either fresh frozen at –80°C or placed into ZBF for fixation. DAI scoring was completed on each mouse after euthanasia using stool collected from the rectum.

### Histopathology

After 72 hours of fixation in ZBF, the colons were cut longitudinally, Swiss-rolled, pinned, and placed in 70% ethanol. Colon was paraffin embedded, sectioned at 4µm thickness, deparaffinized, and stained with hematoxylin and eosin (H&E) by the University of Washington Diabetes Research Center Histology and Imaging Core (HIC). Colonic tissue stained with H&E was imaged using brightfield microscopy on the Keyence BZ-X800 microscope (Keyence Corp. of America, Itasca, IL, United States). A total of 8 random images of intact colon tissue at x 10 magnification were taken from each animal. A modified scoring protocol from Wirtz *et al* was used to numerically score the extent of tissue damage by a blinded pathologist [49]. The score encompasses tissue damage (0 = none, 1 = isolated focal epithelial damage, 2 = mucosal erosions and ulcerations, 3 = complete ulceration/loss of architecture), extent of tissue damage (0 = no sites involved, 1 = 1-24% involved, 2 = 25-50% involved, 3 = >50% involved), degree of inflammatory cell infiltrate (0 = no increase, 1 = increase in lamina propria, 2 = increase in submucosa, 3 = increase through muscularis propria along vascular arcades, 4 = increase in the serosa), and extent of inflammatory cell infiltrate (0 = no increase in inflammatory cells OR no inflammatory cells penetrating into the submucosa, 1 = 1-24% of bowel length involved, 2 = 25-50% of bowel length involved, 3 = >50% of bowel length involved).

### Immunofluorescence

Paraffin embedded colon samples were deparaffinized with xylene and stored in 0.1 M PBS azide. Slides were then subjected to antigen retrieval using heated 10mM tri-sodium citrate with 500 μL Triton-X pH 8. After cooling, slides were permeabilized using 0.2% Triton X-100 in PBS for 25 minutes with three subsequent 0.05% PBS-T washes. Slides were incubated with primary antibodies to claudin-1 (1:500, Abcam, Cambridge, United Kingdom) and E-cadherin (1:500, Cell Signaling, Danvers, MA), for two hours. Sections were washed in 0.05% PBS-T and incubated overnight with Alexa 647-conjugated donkey anti-mouse antibody, Alex 555-conjugated donkey anti-rabbit or anti-chicken antibody, and 488-conjugated donkey anti-rabbit antibody. After 24 hours, slides were washed in 0.05% PBS-T, incubated with DAPI for 20 minutes, washed in 0.05% PBS-T, and incubated with 1X True Black® Lipofuscin autofluorescence quencher in PBS (40X Biotium 23014, Fremont, CA) for 5 minutes. Slides were air dried overnight and cover-slipped with Fluoromount-G^TM^ (Invitrogen 00-4958-02, Carlsbad, CA). Immunofluorescence images were captured using the Keyence BZ-X800 microscope at 20X magnification. Mean fluorescence intensity analysis was performed using open-source ImageJ software building on published protocols [50].

### Mass Spectroscopy

As previously published by our group [35], paraffin-embedded colon samples were deparaffinized with xylene and then washed in 0.1 M phosphate buffered saline (PBS) followed by 3 x in Optima liquid chromatography/mass spectrometry (LC/MS) – grade water (Thermo Fisher Scientific, 7732-18-5, Waltham, MA) and 1 x in 50 mM ammonium bicarbonate pH 7.6; Sigma, 09830, Burlington, MA). Samples were then exposed to ChondroitinaseABC (ChABC) enzyme (500mU/mL in 50mM ammonium bicarbonate, pH 7.6; Sigma, C3667, Burlington, MA) to selectively degrade CS/DS-GAG chains. Slides were incubated in chondroitinase ABC (ChABC) enzyme at 37°C in a Thermo Fischer Scientific MaxQ4000 Orbital Incubator for 24h. Samples were then collected and centrifuged at room temperature to remove tissue debris, dehydrated and concentrated using a Thermo Fisher Scientific SpeedVac Concentrator, then reconstituted to 30 μl in Optima water, with subsequent analysis with liquid chromatography tandem mass spectrometry (LC-MS/MS). Tissue processing steps are outlined in Figure 6A. A triple quadrupole mass spectrometer equipped with electrospray ion source (Waters Xevo TQ-S, Milford, MA) operated in negative mode ionization with a Hypercarb column (2.1 Å∼50mm, 3mm; Thermo Fisher Scientific, Waltham, MA) was used to complete the analysis, and data was acquired and quantified using MassLynx software version 4.1 (Waters, Milford, MA). Peaks were analyzed in the multiple reaction monitoring channels as described in previous publications [35, 51] and a modified peak area normalization function using equimolar CS standard mixes was used to express each CS isomer abundance as a percentage of the total CS-GAG content.

### Statistical Analysis

Data from individual experiments including DAI, spleen weight, colon length, %weight loss, % area staining, H&E damage scores, and CS-E abundance data are shown as dot plots representing data from individual animals. All bar graphs represent mean ± SEM. Student’s t-test was used to compare mean values between two groups, while two-way analysis of variance (ANOVA) was used to compare multiple groups. Calculations were completed using GraphPad Prism 9 (GraphPad Software, CA, United States). Animals were not excluded from the study unless otherwise indicated. All differences were considered statistically significant if p<0.05. Error bars in the figures represent the standard error of the mean. Pathology investigators were blinded to group conditions during histologic damage scoring.

## AUTHOR CONTRIBUTIONS

BD, KF, CB, GM, MS, and JS designed and executed all mice experiments and tissue analysis. BD, CB, and JS performed all mice surgeries and post-op care. JS, CB, TK, SM, JK, KK, and BAP assisted with mouse measurements, performing microscopy, and image analysis. KS assisted with image analysis. CW, PC, CB and GM performed experiments related to optimal Semaglutide dose selection prior to these experiments. MP performed blinded tissue H&E analysis with subsequent disease activity scoring. BD wrote the manuscript as first author with subsequent contributions by KF, CB, GM, MS, JS. All authors reviewed and edited the manuscript prior to submission.

## ACKNOWLEDGEMENTS

The authors thank the University of Washington Diabetes Research Center Histology Imaging Core (P30 DK017047) for their assistance with colon tissue processing, sectioning, and staining (H&E). Additionally, the authors thank The Seattle Children’s Department of Pathology (Gail Deutch) for assisting with imaging H&E slides. This work was also supported by the National Institute of Diabetes and Digestive and Kidney Diseases (NIH-NIDDK) grants DK141910 to KF, DK089056 and DK124238 to GJM, DK083042 and DK101997 to MWS, the Diabetes, Obesity, and Metabolism Training Grant (T32 DK007247) to BD, PPC and CAW, a Nutrition Obesity Research Center Pilot and Feasibility award to KLF (P30 DK035816) at the University of Washington, a Dick and Julia McAbee Endowed Fellowship (to CAW) and a DoD grant (W81XWH2110635 (to JMS). NIH National Institute of Allergy and Infectious Disease (NIH-NIAID) grant DP2AI171150 (KMA).

