## Supplemental Figures for "Central Glucagon-Like Peptide-1 Agonism Ameliorates Colonic Inflammation in a Murine Model of Inflammatory Bowel Disease"

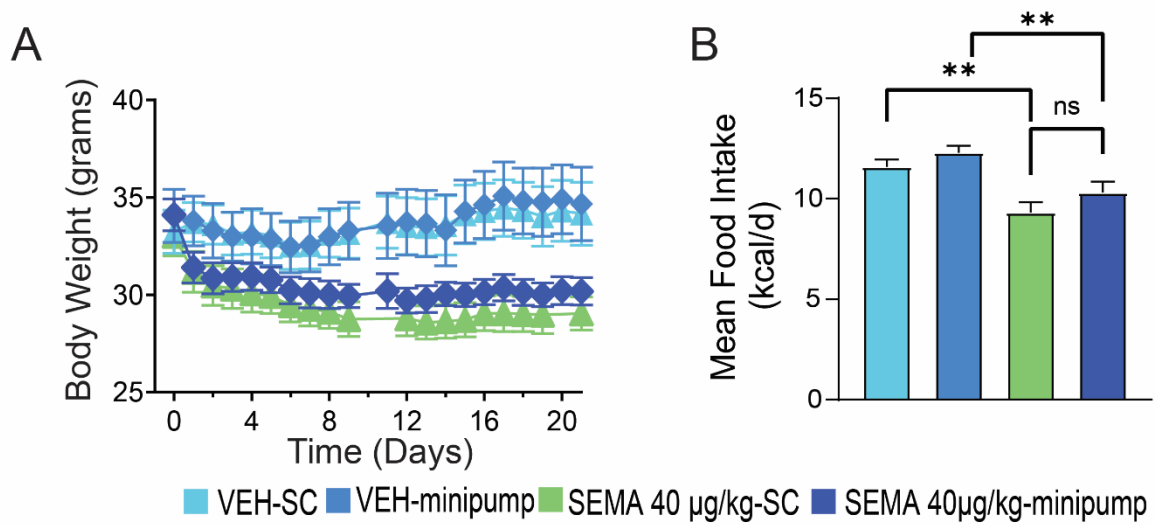

**Supplementary Figure 1. Food intake and body weight of DIO mice receiving SEMA via either osmotic mini-pump or daily subcutaneous injections.** (A) Body weight (grams) and (B) mean food intake (kcal/day) in mice given either SEMA (40ug/kg/day) or VEH via osmotic mini-pump or daily subcutaneous injection. Welch's T-test. SEMA: Semaglutide; VEH: Vehicle; SC: Subcutaneous; DIO: diet-induced obesity. \*\*\*\*P< 0.0001, \*\*\*P< 0.001, \*\*P< 0.01, \*P< 0.05.

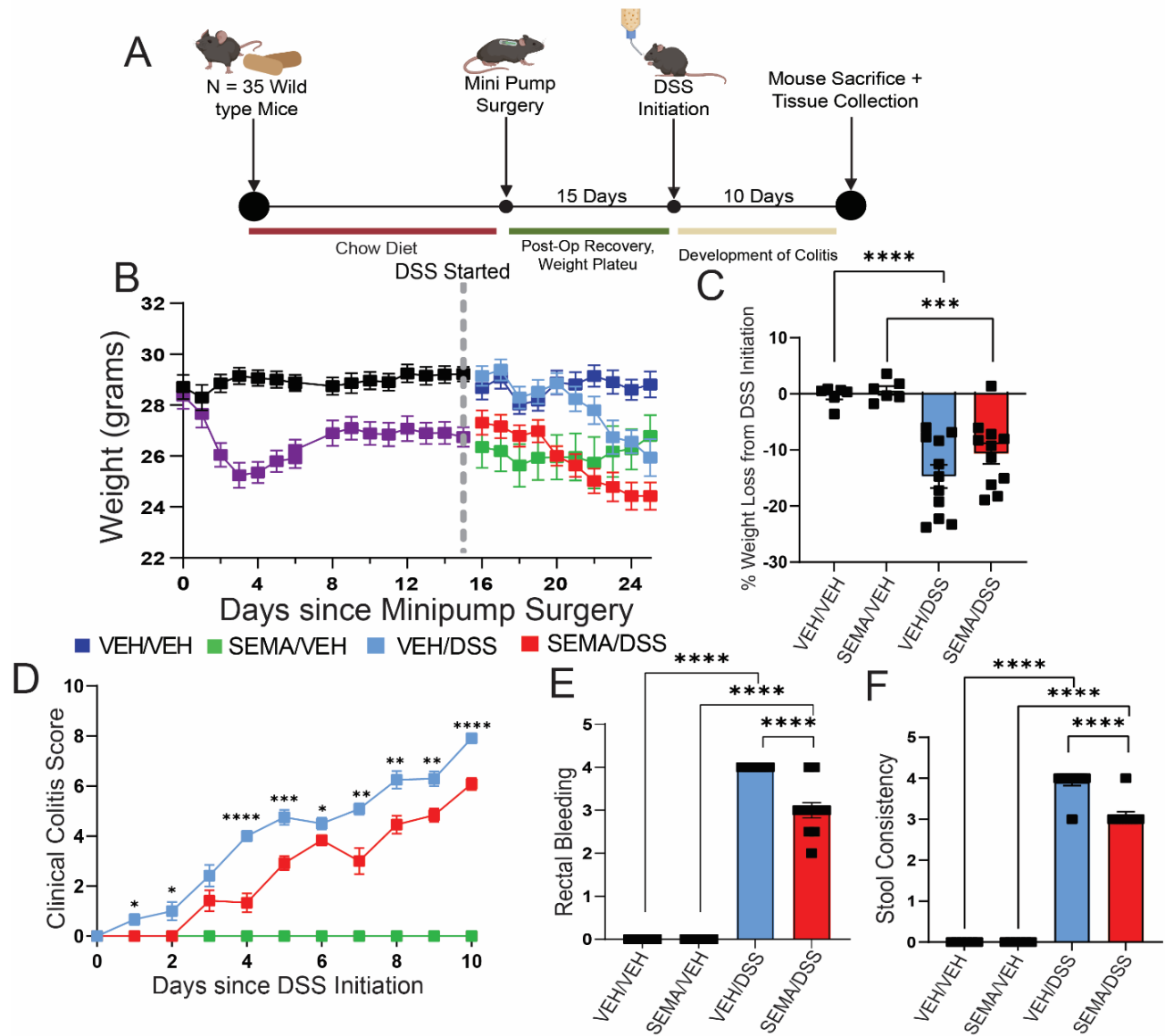

**Supplementary Figure 2. Peripheral SEMA improves clinical outcomes in chow-fed DSS treated mice** (A) Experimental timeline, (B) body weight, (C) clinical colitis scores, (D) total percent weight loss at end of study from time of DSS initiation, (E) rectal bleeding, and (F) stool consistency in chow-fed mice that received continuous subcutaneous infusion of either the GLP-1RA, SEMA (40 μg/kg/d) or its VEH and subsequently received either 2% DSS or VEH in the drinking water for that last 10 days. $n = 11 - 12$ for VEH/DSS and SEMA/DSS groups, $n = 6$ for SEMA/VEH and VEH/VEH groups. Mean ± standard error of the mean for (B) and (C). Non-parametric t-test comparing VEH/DSS and SEMA/DSS in (C). 2-way ANOVA followed by Fisher's post hoc tests for (D-G). SEMA: Semaglutide; VEH: Vehicle; DSS: Dextran Sodium Sulfate; DIO: Diet induced obesity. \*\*\*\*$P < 0.0001$, \*\*\*$P < 0.001$, \*\*$P < 0.01$, \*$P < 0.05$.

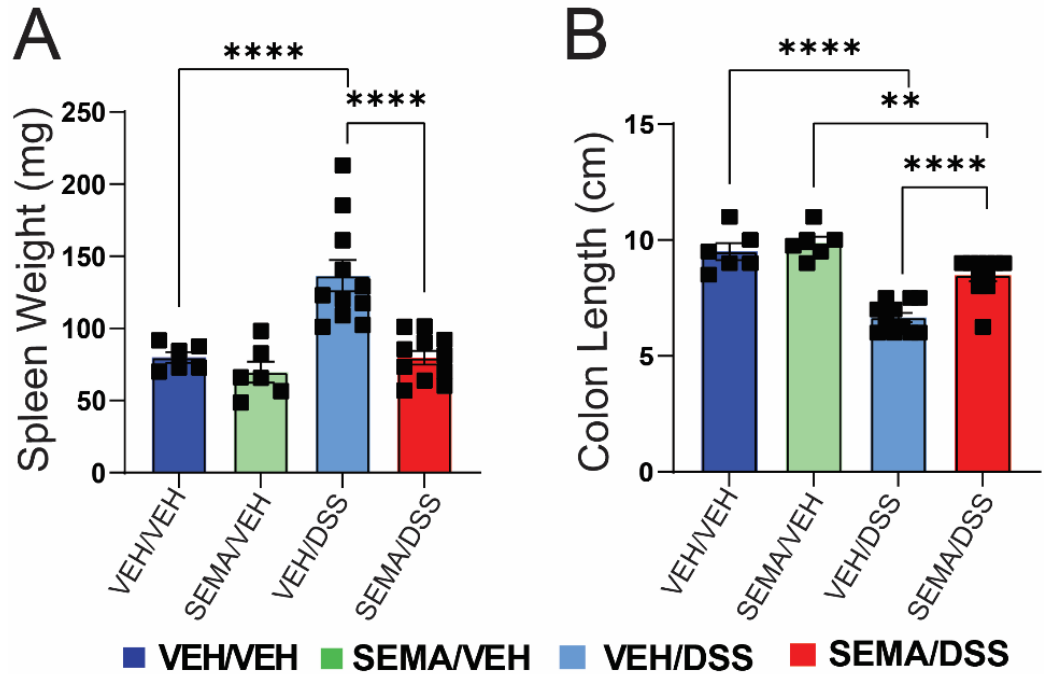

**Supplementary Figure 3. Effect of peripheral SEMA administration on spleen weight and colon length in chow-fed DSS-treated mice.** (A) Spleen weights and (B) colon Length in lean, chow-fed mice that received continuous subcutaneous infusion of either the GLP-1RA, SEMA (40 μg/kg/d) or its VEH and subsequently received either 2% DSS or VEH in the drinking water for that last 10 days. $n = 11 - 12$ for VEH/DSS and SEMA/DSS groups, $n = 5 - 6$ for SEMA/VEH and VEH/VEH groups. 2-way ANOVA followed by Fisher's post hoc tests. SEMA: Semaglutide; VEH: Vehicle; DSS: Dextran Sodium Sulfate; DIO: Diet induced obesity; H&E: Hematoxylin and Eosin. \*\*\*\*$P < 0.0001$, \*\*\*$P < 0.001$, \*\*$P < 0.01$, \*$P < 0.05$.

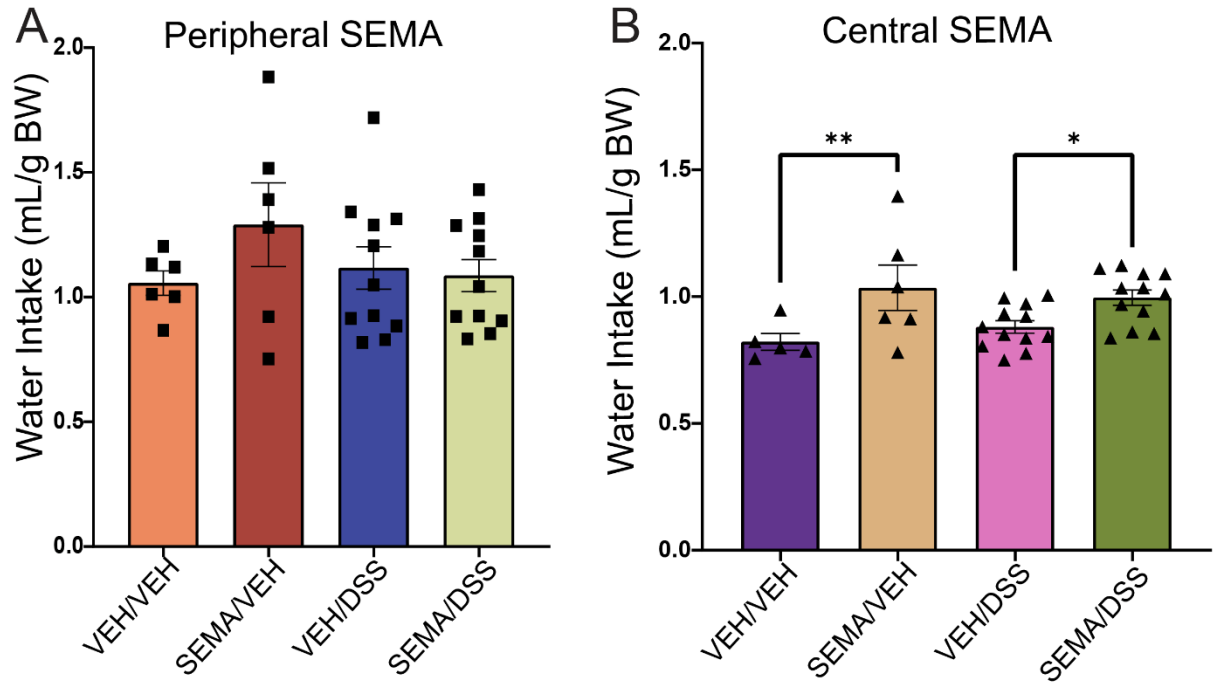

**Supplementary Figure 4. Differences in cumulative water intake (adjusted for body mass) do not associate with disease severity in DSS-treated animals.** Cumulative water intake in (A) peripheral (40 μg/kg/d) and (B) central (4 μg/kg/d) or its VEH during DSS treatment normalized to body weight (g). $n = 11 - 12$ for VEH/DSS and SEMA/DSS groups, $n = 5 - 6$ for SEMA/VEH and VEH/VEH groups. 2-way ANOVA followed by Fisher's post hoc tests. SEMA: Semaglutide; VEH: Vehicle; DSS: Dextran Sodium Sulfate; DIO: Diet induced obesity; \*\*\*\*$P < 0.0001$, \*\*\*$P < 0.001$, \*\*$P < 0.01$, \*$P < 0.05$.

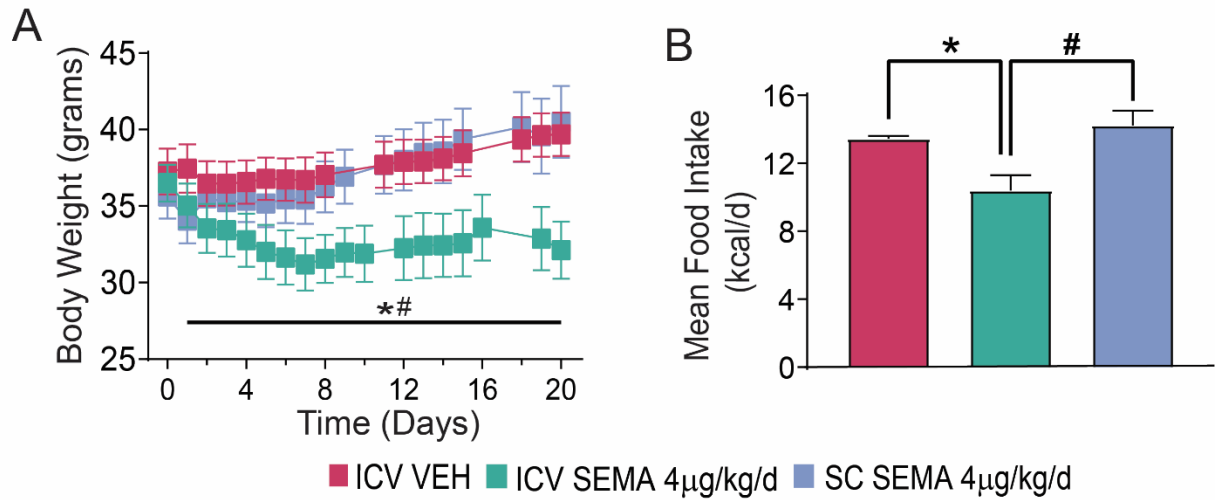

**Supplementary Figure 5. Lack of effect of low-dose SEMA on food intake and body weight when given peripherally to DIO mice. (A)** Body weight (grams) and **(B)** mean food intake (kcal/day) in mice given either SEMA (4ug/kg/day, the dose used for icv infusion) or VEH via peripheral osmotic mini-pump or an osmotic mini-pump attached to an ICV cannula. Welch's T-test. SEMA: Semaglutide; VEH: Vehicle; SC: Subcutaneous; DIO: diet-induced obesity. \*\*\*\*P< 0.0001, \*\*\*P< 0.001, \*\*P< 0.01, \*P< 0.05.

| % CS/DS Isomer<br>Colon<br>mean ± SEM | VEH/VEH | SEMA/VEH | VEH/DSS | SEMA/DSS | p-value |  |  |  |
| --- | --- | --- | --- | --- | --- | --- | --- | --- |
|  |  |  |  |  | VEH/VEH vs.<br>SEMA/VEH | VEH/VEH<br>vs.<br>VEH/DSS | VEH/VEH vs.<br>SEMA/DSS | VEH/DSS<br>vs.<br>SEMA/DSS |
| **CS-A (4S)** | 76.5 ± 0.8 | 76.4 ± 0.9 | 76.5 ± 0.4 | 78.2 ± 0.3 | 0.9 | 0.9 | 0.2 | 0.1 |
| **CS-O (0S)** | 7.7 ± 0.3 | 7.8 ± 0.1 | 7.0 ± 0.2 | 6.5 ± 0.2 | 0.9 | 0.2 | 0.009<br>** | 0.4 |
| **CS-E (4S6S)** | 3.7 ± 0.3 | 4.1 ± 0.4 | 6.5 ± 0.3 | 4.8 ± 0.2 | 0.8 | <0.0001<br>**** | 0.1 | <0.0001<br>**** |
| **DS (2S4S)** | 7.1 ± 0.3 | 6.9 ± 0.6 | 6.8 ± 0.3 | 6.5 ± 0.3 | 0.9 | 0.9 | 0.7 | 0.9 |
| **CS-C (6S)** | 5.1 ± 0.2 | 4.8 ± 0.3 | 3.4 ± 0.1 | 4.1 ± 0.1 | 0.8 | <0.0001<br>**** | 0.003<br>** | 0.01<br>* |
| **Avg. # sulfates**<br>**per CS/DS** | 1.03 ± 0.006 | 1.03 ± 0.008 | 1.06 ± 0.007 | 1.05 ± 0.004 | 0.9 | 0.01<br>* | 0.3 | 0.2 |

**Supplemental Table 1. CS/DS isomers in the colon with peripheral semaglutide administration in a DSS colitis model.**

Relative abundance of colonic CS/DS isomers in diet-induced obese mice with DSS-induced colitis after receiving vehicle or intraperitoneal semaglutide.
